# The fungal-specific Hda2 and Hda3 proteins regulate morphological switches in the human fungal pathogen *Candida albicans*

**DOI:** 10.1101/340364

**Authors:** Misty R. Peterson, Robert Jordan Price, Sarah Gourlay, Alisha May, Jennifer Tullet, Alessia Buscaino

## Abstract

The human fungal pathogen *Candida albicans* is responsible for millions of infections annually. Due to the few available anti-fungal drugs and the increasing incidence of drug resistance, the number of *C. albicans* infections is dramatically increasing. Morphological switches, such as the white-opaque switch and the yeast-hyphae switch, are key for the development of *C. albicans* pathogenic traits. Lysine deacetylases are emerging as important regulators of morphological switches. Yet, targeting lysine deacetylases for drug development is problematic due to the high homology between the fungal and human proteins that could result in toxicity. Here we provide evidence that the fungal specific proteins Hda2 and Hda3 interact with the lysine deacetylase Hda1. By combining phenotypic analyses with genome-wide transcriptome analyses, we demonstrate that Hda2 and Hda3 control *C. albicans* morphological switches. Under nutrient-rich conditions, deletion of *HDA2* or *HDA3* leads to moderate overexpression of the master regulator of white-opaque switching *WOR1* and increase switching frequency. Under hyphae inducing conditions, deletion of *HDA2* and *HDA3* block hyphae development. However, deletion of *HDA2* and *HDA3* does not affect hyphae-formation and virulence *in vivo*. We propose that Hda2 and Hda3 are good targets for the development of anti-fungal drugs to be used in combination therapy.

## INTRODUCTION

Fungal pathogens are a leading cause of human mortality causing over 1.5 million deaths per year (1). *C. albicans* is a commensal organism that colonises the mouth, gastrointestinal and reproductive tract of healthy individuals without causing any harm. Still, *Candida albicans* is also the most common human fungal pathogen and the principal causal agent of mycotic death (2). This is because, in immune-compromised patients, *C. albicans* can invade vital organs and cause serious, life-threating systemic infections associated with a mortality rate up to 70 *%* (2). The ability to transition between different morphological forms in response to changing environments is a key virulence trait in *C. albicans*.

For example, *C. albicans* cells can reversibly switch between white and opaque forms (3). White and opaque cells are genetically identical, yet they differ in cellular morphology, colony shape, gene expression profile and mating behaviour (4). In addition, white cells are more virulent in a murine model of systemic infection (5, 6) whereas opaque cells preferentially colonise the skin (7). White-opaque switching is under the control of the master regulator Wor1, a transcription factor whose expression is necessary and sufficient for opaque cell formation (8–11). Stochastic increases in Wor1 levels drives the transition from the white to the opaque phase. Furthermore, Wor1 expression produces a direct positive feedback loop by binding its own promoter and turning on its own expression (8, 9, 11). Switching is also regulated by the mating type locus as opaque formation occurs predominantly in a or α cells (12).

*C. albicans* virulence also depends on its ability to convert between yeast and hyphal morphology: yeast cells are critical for colonisation, early infection and dissemination, while hyphal growth is responsible for tissue invasion and chronic infections (13). Hyphal morphogenesis is coupled with virulence, as several of the genes, that are specifically expressed in hyphae, encode virulence factors (14–17). Hyphal morphogenesis is a complex and highly orchestrated process, and *C. albicans* uses multiple redundant pathways to integrate host signals and promote hyphae development. Indeed, *C. albicans* filamentation can be induced by many environmental cues such mammalian serum, body temperature, hypoxia and CO_2_ concentration which reflects the variety of signals sensed by the fungus in the different microenvironments encountered in the host. In yeast cells, hyphae morphogenesis is inhibited by the DNA-binding repressor Nrg1 that, together with Tup1, blocks expression of a subset of filament-specific genes (18). Upon hyphal induction, filamentous growth is promoted by Efg1 and Flo8, two transcription regulators essential for hyphal development and virulence [14–16]. During the initiation phase of hyphae morphogenesis, Nrg1-mediated repression is cleared via *NRG1* transcriptional repression and Nrg1 protein degradation. During hyphae maintenance, chromatin remodelling of hyphae promoters prevents Nrg1 binding despite its increased protein levels (19).

Due to the close evolutionary relationship between fungi and the human host, effective treatment for *C. albicans* infections is hindered by the limited number of sufficiently divergent potential drug targets. There are only three classes of antifungal drugs effective for the treatment of systemic fungal infections and their clinical utility is limited by the rapid emergence of drug resistance (1, 20). Hence, there is an immediate and urgent need to develop alternative treatments. A potential strategy is to target *C. albicans* morphological plasticity.

Lysine deacetylases (KDACs, also known as HDACs) act as global regulators of gene expression by catalysing the removal of acetyl functional groups from the lysine residues of histones and non-histone proteins (21). KDACs can favour transcriptional repression by deacetylating lysine residues on histone tails allowing chromatin compaction and/or preventing binding of bromodomain-containing transcriptional activators (21). KDACs can also activate transcription by deacetylation of non-histone proteins (22). As a consequence, deletion or inhibition of KDACs often results in the upregulation and downregulation of an approximately equivalent number of genes (23). KDACs are highly conserved across eukaryotes and can be phylogenetically divided into three main classes: the Rpd3 and Hos2-like (class I) enzymes, the Hda1-like (class II) enzymes and the Sir2-like (class III) enzymes. The class I and class II enzymes are related, sharing a conserved central enzymatic domain. The class III enzymes are nicotinamide adenine dinucleotide (NAD) dependent. KDACs lack intrinsic DNA-binding activity and are recruited to target genes via incorporation into large multiprotein complexes or direct association with transcriptional activators and repressors (24).

In *C. albicans*, the lysine deacetylase Hda1 (class II) is an important regulator of morphological switches. Hda1 controls white-opaque switching as deletion of the *HDA1* gene increases switching rates from white to opaque (25, 26). In response to serum, N-acetylglucosamine or nutrient limitation, Hda1 also controls the yeast to hyphae switch by deacetylating Yng2, a component of the histone acetyltransferase NuA4 complex, blocking maintenance of hyphal growth (19). However, the Hda1 pathway is not required for hyphae elongation in hypoxia or in the presence of elevated CO_2_ because of the presence of redundant pathways (27, 28). As a result, the Hda1-mediated hyphae maintenance pathway contributes, but it is not absolutely required, for virulence *in vivo* (27). These results suggest that Hda1 is a good target for antifungal drugs development to be used in combination therapies.

KDACs are promising druggable targets: one KDAC inhibitor is currently used for cancer treatment and several other KDAC inhibitors are in clinical trials (29, 30). However effective targeting of Hda1 for anti-fungal drug development is impaired by the high sequence similarities between Hda1 and its human orthologs and consequently a likeliness for high toxicity.

In *Saccharomyces cerevisiae*, Hda1 assembles with two non-catalytic subunits, Hda2 and Hda3, essential for Hda1 deacetylation activity both *in vivo* and *in vitro* (31). Interestingly, no metazoan homologous of Hda2 and Hda3 have been identified. Hda2 and Hda3 are similar in sequence and share a similar protein organisation with an N-terminal DNA binding domain (DBD) and a C-terminal coil-coil domain (CCD). The DBD domain, similar in structure to the helicase domain of the SWI-SNF chromatin remodellers, is sufficient to bind DNA *in vitro*. The CCD domains act as a scaffold for the assembly of the Hda1 complex (32).

Here for the first time, we analyse the role of *C. albicans* Hda2 and Hda3. We demonstrate that the Hda1 complex is conserved in *C. albicans* as Hda2 and Hda3 interact with Hda1 *in vivo*. Our analyses demonstrated that, in yeast-inducing conditions, deletion of *HDA2* and *HDA3* leads to transcriptional upregulation of *WOR1* and that Hda2 and Hda3 inhibit white and opaque switching. In contrast, under hyphae-inducing conditions, Hda2 and Hda3 regulate the yeast to hyphae switch. This functional rewiring is linked to a reduced expression of components of the Hda1 complex. Our study demonstrates that Hda2 and Hda3 control key morphological switches in *C. albicans*. Therefore, we propose that Hda2 and Hda3 are attractive targets for the development of novel antifungal drugs.

## RESULTS

### The Hda1 complex is conserved in *C. albicans*

*S. cerevisiae* Hda2 and Hda3 are critical components of the Hda1 complex as they are required for the Hda1-mediated histone deacetylation activity (31). BLAST analyses reveal that the *C. albicans* genome contains two genes encoding proteins with homology to *S. cerevisiae* Hda2 (C3_03670W) and Hda3 (CR_09490W_A). Structural alignments predict that Hda2 and Hda3 have a similar protein organisation with an N-terminal DNA binding domain (DBD) and a C-terminal coil-coil domain (CCD) (Fig 1A and 1B). To explore the potential functional relationship between *C. albicans* Hda2, Hda3 and Hda1, we assessed whether we could detect a physical interaction between these proteins. To this end, we generated strains expressing at the endogenous locus an epitope-tagged Hda1 protein (Hda1-HA) together with either Hda2-GFP or Hda3-GFP. Western analyses show that the Hda1-HA and Hda2-GFP tagged proteins are expressed at high levels while we could not detect Hda3-GFP in whole cell extract indicating that the protein is expressed at low levels (Fig 1C). Immuno-precipitation (Ip) of Hda1-HA with a highly specific anti-HA antibody demonstrates that Hda2 strongly interacts with Hda1. We could also detect an interaction between Hda1 and Hda3, despite the low level of expression of Hda3 (Fig 1C). Thus, Hda2 and Hda3 physically interact with Hda1. To delineate the function of *C. albicans* Hda2 and Hda3, we generated homozygous deletions mutants for *HDA2* (*hda2 Δ/Δ*) and *HDA3* (*hda3 Δ/Δ*) genes using a recyclable Clox system (Fig 1D) (33). The Clox system allows for the generation of a homozygous deletion mutant lacking any marker genes and therefore genetically identical to the parental strain except for the deleted gene. This allows for the direct comparison of phenotypes between mutant and parental strains. Growth analyses reveal that Hda2 and Hda3 are not required for survival and fitness as the *hda2 Δ/Δ* and *hda3 Δ/Δ* strains grow similarly to the wild-type (WT) control both on solid and in liquid media (Fig 1E and 1F). Therefore, *C. albicans* Hda2 and Hda3 are *bonafide* components of the Hda1 complex that are not required for cell fitness and survival under optimal growth conditions.

**Figure 1.**
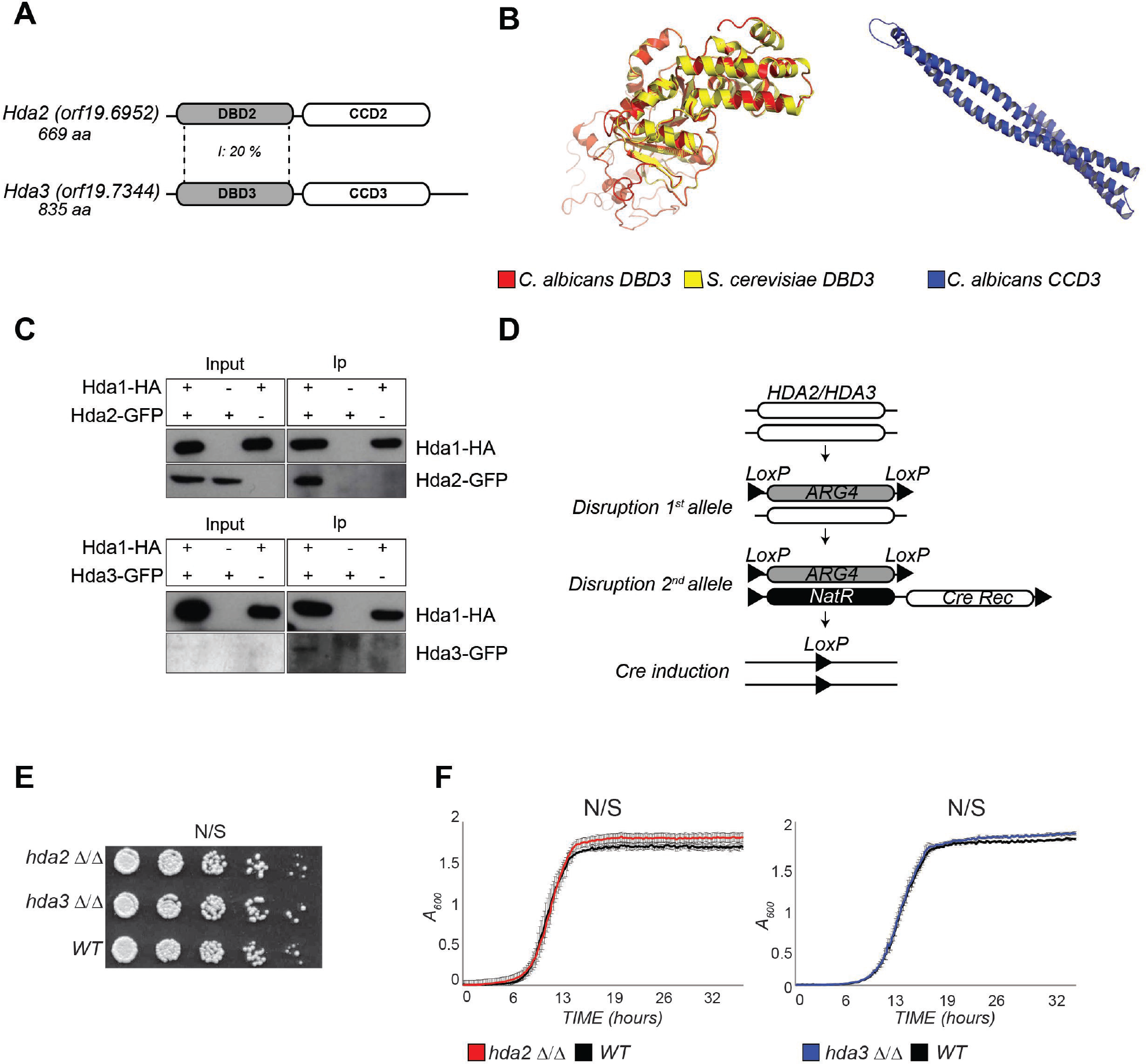
The Hda1 complex is conserved in *C. albicans*. **(A)** Domain organisation of *C. albicans* Hda2 and Hda3 proteins **(B)** *Left:* Structural alignment of *C. albicans* DBD3 domain (red) with *S. cerevisiae* DBD3 domain (yellow); *Right:* Structural modelling of *C. albicans* CCD3 **(C)** Co-Immunoprecipitation of Hda1 with Hda2 and Hda3. Hda1-HA Immunoprecipitation (Ip) analysed with anti-HA or with anti-GFP to detect Hda2 and Hda3. **(D)** Schematic of *Clox* gene disruption strategy used to construct *hda2*Δ/Δ and *hda3*Δ/Δ mutants. **(E)** Serial dilution assay of WT, *hda2*Δ/Δ and *hda3*Δ/Δ mutants on solid YPD media at 30 °C. **(F)** Growth curves of WT, *hda2*Δ/Δ and *hda3*Δ/Δ isolates in YPD liquid media at 30 °C.

### Global gene expression changes in the absence of Hda2 and Hda3

To gain insights about the function of Hda2 and Hda3 in conditions stimulating yeast growth, we analysed changes in gene expression by strand-specific RNA-sequencing (RNA-seq) upon deletion of the *HDA2* and *HDA3* genes. As a comparison, we also performed RNA-seq on strains with the *HDA1* gene deleted. Deletion of *HDA2* results in 577 genes significantly differentially regulated (348 upregulated and 229 downregulated, q < 0.05; Dataset S1). These gene expression changes are highly similar to the ones observed in *hda1 Δ/Δ* (Pearson correlation coefficient r = 0.7, Dataset S1) (Fig 2A). Deletion of the *HDA3* gene has a less profound effect on gene expression as only 78 genes are significantly differentially expressed (45 upregulated and 33 downregulated, q < 0.05; Dataset S1). Despite this difference, changes in *hda3 Δ/Δ* cells correlate with *hda1 Δ/Δ* and *hda2 Δ/Δ* gene expression changes (Pearson correlation coefficient r = 0.66 and 0.46, respectively) (Fig 2A).

**Figure 2.**
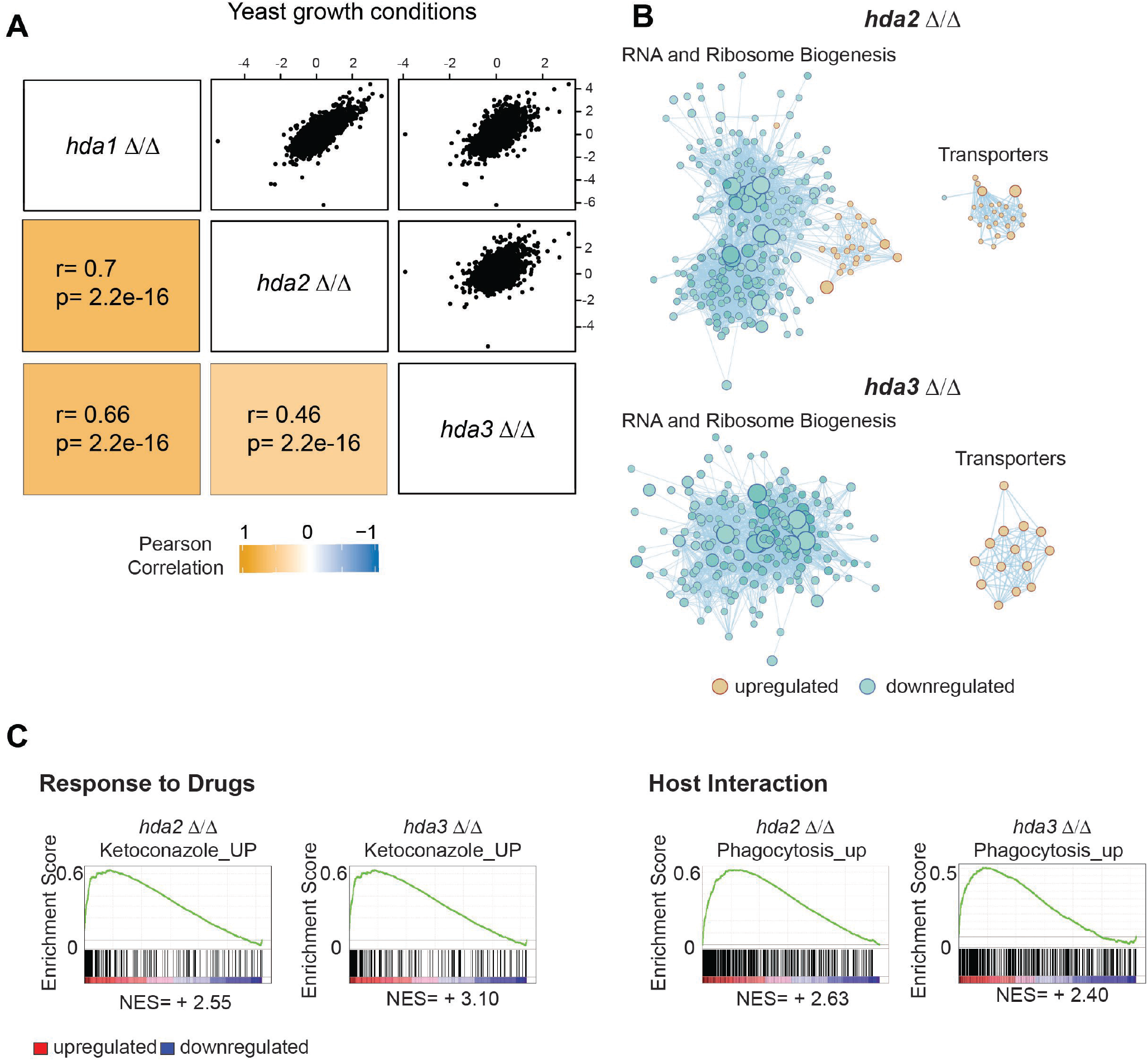
Global gene expression changes in the absence of Hda2 and Hda3. **(A)** Pearson correlation matrix of gene expression changes observed in *hda1 Δ/Δ, hda2 Δ/Δ* and *hda3* Δ/Δ grown in YPD at 30 °C. r = Pearson correlation coefficient. p = p-value. **(B)** The network of functional groups of genes regulated by Hda2 and Hda3 constructed by GSEA and Enrichment Map. Blue circles represent down regulated while orange circles depict upregulated gene sets which are linked in the network by grey lines. The diameter of the circles varies based upon the number of transcripts within each set. **(C)** Example enrichment plots for selected genes sets differentially expressed in *hda2* Δ/Δ (left) and *hda3* Δ/Δ (right). Ketoconazole_up: set of genes upregulated in *C. albicans* cells grown in the presence of ketoconazole (59); Phagocytosis_up: gene set upregulated following engulfment by primary Bone Marrow Derived Macrophages (60). The x-axis shows genes ranked according to their expression in the mutants from up-regulated (left) to down-regulated (right) genes. Black vertical lines mark individual genes. The cumulative value of the enrichment score (y-axis) is represented by the green line. A positive normalised enrichment score (NES) indicates enrichment in the up-regulated group of genes in *hda2* Δ/Δ and *hda3* Δ/Δ.

To reveal the cellular pathways regulated by Hda2 and Hda3 in *C. albicans*, we performed Gene Set Enrichment Analysis (GSEA) of the RNA-seq datasets (34). To this end, transcript profiles of *hda2 Δ/Δ* or *hda3 Δ/Δ* isolates were ranked according to their differential expression and this list was compared to gene sets identified in other experimental analyses (35) and (Sellam et al, personal communications). This allows the identification of statistically significant gene sets enriched in the top (up-regulated) or the bottom (down-regulated genes) of the ranked list (34). The network of similar gene sets was visualized using Cytoscape where nodes represent gene sets, and lines connect nodes sharing a significant number of genes (36).

GSEA detected enrichment for rRNA and ribosome biogenesis genes and genes involved in transport (Fig 2B). Enrichment was also found in gene sets important for virulence-promoting function in *C. albicans*. This includes genes differentially expressed during *C. albicans-host* interactions with mouse macrophages and in response to drugs (Fig 2C).

Modulation of several gene sets enriched in *hda2 Δ/Δ* and/or *hda3 Δ/Δ* mutants, for example, down-regulation of genes required for protein synthesis, are part of stress response in *C. albicans* (37). Accordingly, we subjected the *hda2 Δ/Δ* and *hda3 Δ/Δ* mutants to a phenotypic analysis by applying a set of different stress conditions (38). For most conditions, we did not observe any difference between the WT and the mutant strains. However, lack of Hda2 or Hda3 leads to sensitivity to copper, sodium chloride and sodium nitroprusside (SNP) with salicylhydroxamic acid (SHAM). Strains mutant in these proteins also show resistance to rapamycin (Fig 3) indicating a role for Hda2 and Hda3 in specific stress responses.

**Figure 3.**
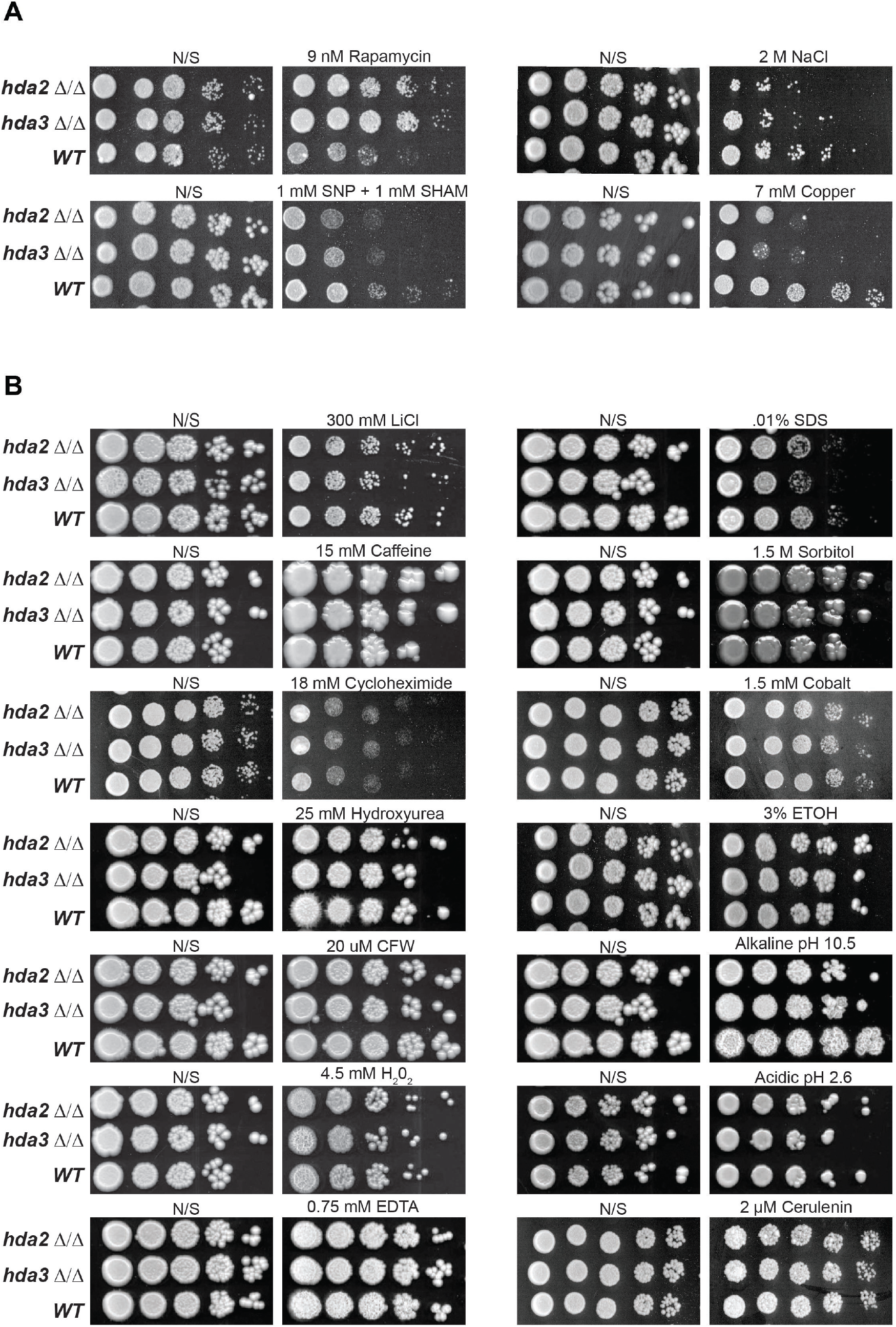
Phenotyping of *hda2* Δ/Δ and *hda3* Δ/Δ strains. Serial dilution assay showing growth of *hda2* Δ/Δ and *hda3* Δ/Δ mutants on solid YPD media with additives as indicated and incubation at 30 °C for 2 - 4 days. **(A)** Additives affecting *hda2* Δ/Δ and *hda3* Δ/Δ growth relative to wildtype strain. Mutants are resistant to rapamycin and sensitive to SNP and SHAM, sodium chloride and copper sulphate. **(B)** Conditions eliciting normal growth of mutant strains relative to matching wildtype.

### Hda2 and Hda3 inhibit white-opaque switching

The GSEA analysis identifies white-opaque switching as a process potentially regulated by Hda2 and Hda3 as genes upregulated in the opaque state are also upregulated upon deletion of *HDA2* and *HDA3* (Fig 4A and 4B). Wor1 is the master regulator of white-opaque phenotypic switching and a stochastic increase in Wor1 levels drives the transition from the white to the opaque phase (8, 9, 11). Therefore, we asked whether *WOR1* expression levels were increased in *hda2 Δ/Δ* and *hda3 Δ/Δ* isolates compared to WT cells.

**Figure 4.**
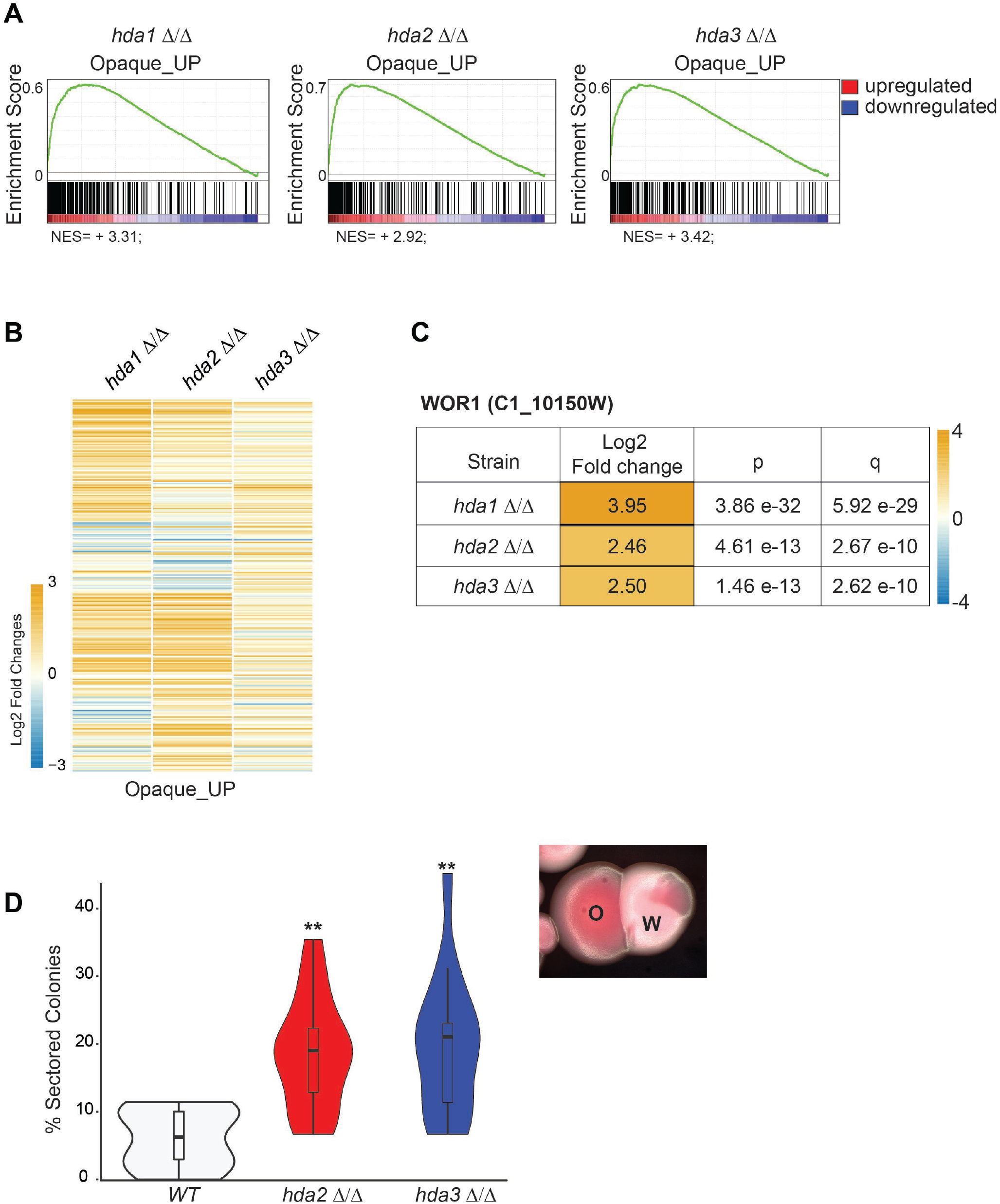
Hda2 and Hda3 inhibit white-opaque switching. **(A)** Enrichment plots for genes differentially expressed in *hda1* Δ/Δ, *hda2* Δ/Δ and *hda3* Δ/Δ in comparison to genes upregulated in opaque cells (38). The x-axis shows genes ranked according to their expression in the mutants from up-regulated (left) to down-regulated (right) genes. Black vertical lines mark individual genes. The cumulative value of the enrichment score (y-axis) is represented by the green line. A positive normalised enrichment score (NES) indicates enrichment in the up-regulated group of genes in *hda1* Δ/Δ, *hda2* Δ/Δ and *hda3* Δ/Δ. **(B)** Heat map depicting the log2 fold change in *hda1* Δ/Δ, *hda2* Δ/Δ and *hda3* Δ/Δ isolates compared to WT for the Opaque_up gene set. **(C)** Log2 fold change values and significance for *WOR1* gene expression in *hda1* Δ/Δ, *hda2* Δ/Δ and *hda3* Δ/Δ compared to WT. **(D)** *Left:* Percentage of sector colonies in WT, *hda2* Δ/Δ and *hda3* Δ/Δ isolates. ** = p value ≤ .01. *Right:* Representative image of cells grown on Phloxine B agar. O = opaque; W = white.

The RNA sequencing analysis demonstrates that deletion of *HDA2* or *HDA3* leads to transcriptional upregulation of *WOR1* (Fig 4C). This result suggests that, in WT cells, Hda2 and Hda3 repress *WOR1* transcription and inhibit white-opaque switching. To test this hypothesis, we generated *MTL a/a* homozygous *hda2 Δ/Δ* and *hda3 Δ/Δ* mutants and measured the frequency of white to opaque conversion using quantitative switching assays. Briefly, cells were plated on Phloxine B plates and the frequency of opaque colonies or colonies containing at least one opaque sector was scored. This analysis demonstrates that deletion of both *HDA2* and *HDA3* increases the frequency of white-opaque switching (Fig 4D). Thus, Hda2 and Hda3 control white-opaque switching, a process that is linked to *WOR1* overexpression.

### Hda2 and Hda3 contribute to filamentous growth but not virulence

To investigate the role of Hda2 and Hda3 under hyphae-inducing conditions, we performed RNA-seq analyses of WT, *hda2 Δ/Δ* and *hda3 Δ/Δ* strains grown in RPMI, a medium which mimics human physiological conditions and therefore strongly induces hyphal growth. As a control, RNA-seq analyses were also performed in *hda1 Δ/Δ* strain. GSEA analysis of the gene expression changes of WT cells grown in yeast-inducing conditions (YPD) and hyphae-inducing conditions (RPMI) confirmed the validity of our experimental approach as genes reported to be upregulated in hyphae compared to yeast are less expressed in YPD compared to RPMI while genes with a yeast-specific expression are expressed at higher levels in YPD than RPMI (Fig 5A).

**Figure 5.**
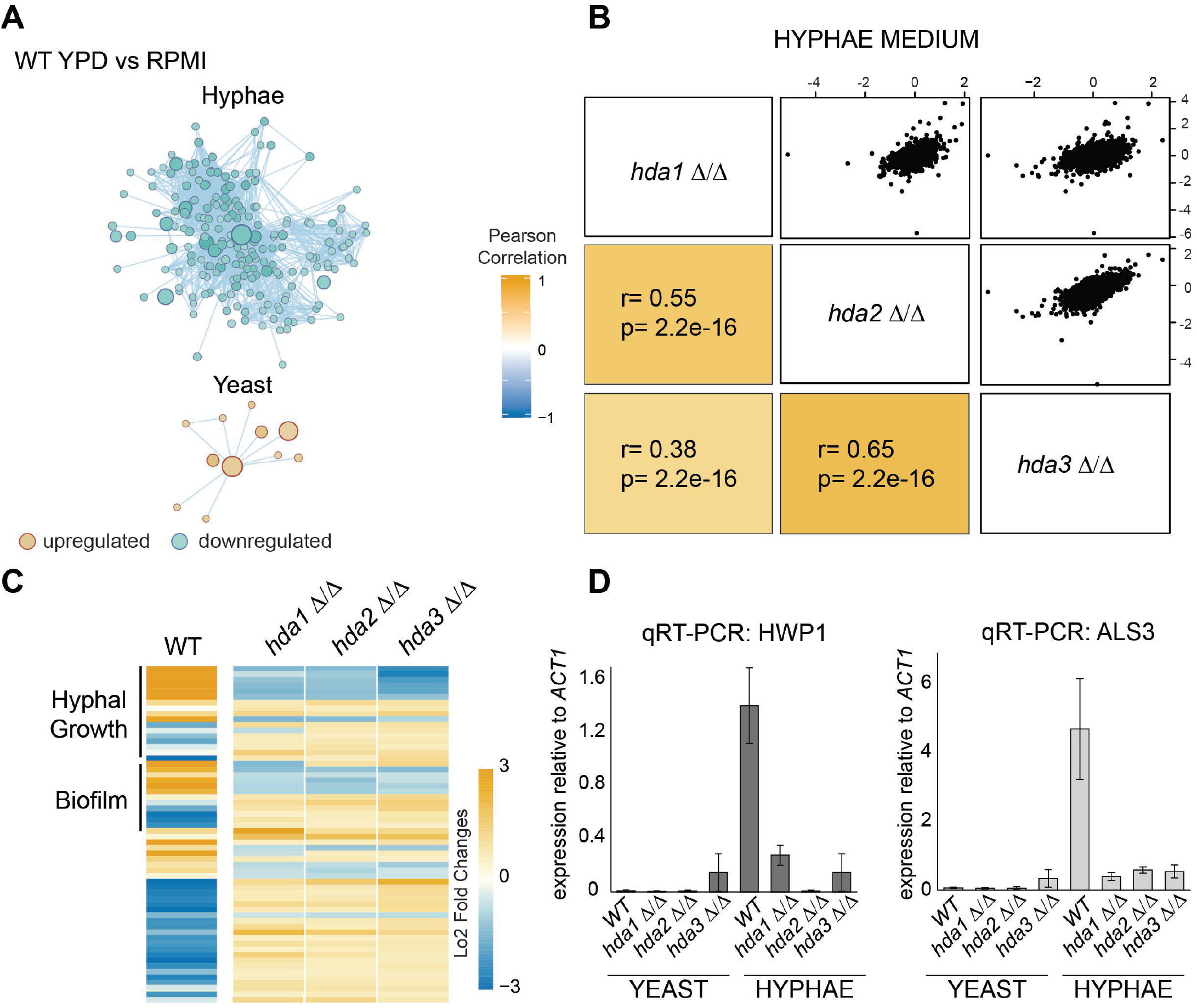
Gene expression profiling in hyphae-inducing conditions in WT, *hda1* Δ/Δ, *hda2* Δ/Δ and *hda3* Δ/Δ isolates. **(A)** The network representing changes in gene expression in WT cells grown in yeast-inducing conditions (YPD) versus hyphae-inducing conditions (RPMI) constructed by GSEA and Enrichment Map. Blue circles represent down regulated gene sets, while orange depicts upregulated gene sets which are linked in the network by grey lines that indicate function. The diameter of the circles varies based upon the number of transcripts within each set. **(B)** Pearson correlation matrix of gene expression changes observed in *hda1 Δ/Δ, hda2 Δ/Δ* and *hda3* Δ/Δ grown in in hyphae growth media (RPMI) at 37 °C. r = Pearson correlation coefficient. p = p-value. **(C)** *Left:* Heat map depicting the log2 fold change in WT cells grown in hyphae-inducing conditions (RPMI at 37 °C) versus WT cells grown in yeast-inducing conditions (YPD at 30 °C). Gene known to be involved in hyphae formation or biofilm formation are indicated. *Right:* Log2 fold changes of hyphae-induced and repressed-genes in *hda1 Δ/Δ, hda2 Δ/Δ* and *hda3 Δ/Δ* isolates. Cell were grown in RPMI at 37 °C. **(D)** Quantitative reverse transcriptase PCR (qRT-PCR) analyses to measure *HWP1* and *ALS3* transcript levels in WT, *hda1 Δ/Δ, hda2 Δ/Δ* and *hda3 Δ/Δ* isolates grown in yeast (YPD at 30 °C) or hyphae (RPMI at 37 °C) inducing conditions. Transcripts levels are visualized relative to *ACT1* transcript levels. Error bars in each panel: standard deviation of three biological replicates.

In hyphae-inducing conditions, deletions of *HDA2* and *HDA3* genes result in 350 and 484 significantly differentially expressed genes, respectively (q< 0.05; Dataset S1). Gene expression changes in *hda2 Δ/Δ* and *hda3 Δ/Δ* correlate with gene expression changes observed in *hda1 Δ/Δ* although less profoundly than in yeast-inducing conditions (Pearson correlation coefficient, r = 0.55 and 0.38, respectively; Fig 5B). The RNA-seq analysis reveals that filamentation is the major pathway misexpressed in the deletion mutants as hyphae-responsive genes are differentially expressed in *hda2 Δ/Δ* and *hda3 Δ/Δ* isolates compared to WT cells (Fig 5C). qRT-PCR analyses of two hyphal growth markers, *HWP1* and *ALS3*, confirmed the RNA-seq results as, in hyphae inducing conditions, expression of *HWP1* and *ALS3* is downregulated in *hda2 Δ/Δ* and *hda3 Δ/Δ* but not in WT cells (Fig 5D).

Collectively, our results suggest that Hda2 and Hda3 are important for hyphal growth. To test this hypothesis, we assessed the morphology of WT, *hda2 Δ/Δ* and *hda3 Δ/Δ* strains upon growth in two different hyphae-inducing media (RPMI and Spider). While WT cells form hyphae efficiently, hyphal growth is defective in *hda2 Δ/Δ* and *hda3 Δ/Δ* cells on both solid and in liquid media (Fig 6A and B). This phenotype was rescued by reintroduction of the *HDA3* gene (Fig 6C). Therefore, Hda2 and Hda3 are important regulators of the yeast to hyphae switch.

**Figure 6.**
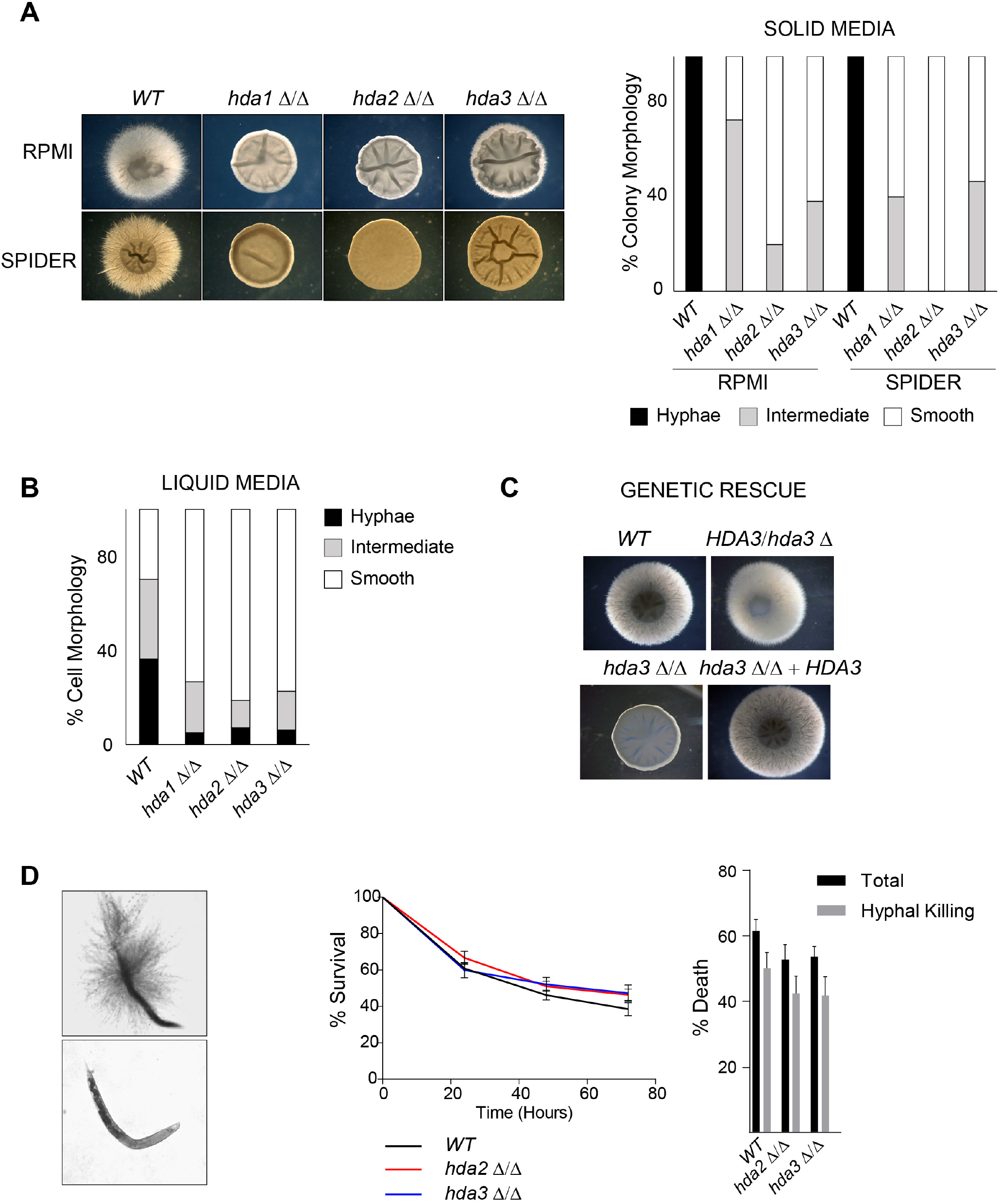
Hda2 and Hda3 contribute to filamentation growth but not virulence. **(A)** *Left:* Representative images of colony morphology of WT, *hda1 Δ/Δ, hda2 Δ/Δ* and *hda3 Δ/Δ* grown on hyphal inducing Spider and RPMI agar at 37 °C. *Right:* Quantification of colony morphologies. **(B)** Quantification of cellular morphologies of WT, *hda1 Δ/Δ, hda2 Δ/Δ* and *hda3 Δ/Δ* grown in liquid RPMI media at 37 °C. **(C)** Rescue experiment of colony morphology upon genomic integration of the *HDA3* gene in the *hda3 Δ/Δ* mutant background (*hda3 Δ/Δ+ HDA3)*. WT, heterozygous (*HDA3/ hda3* Δ) and homozygous *hda3 Δ/Δ* isolates were included as a control. Cell were grown on hyphal inducing RPMI agar at 37 °C. **(D)** *Left:* Representative images of dead *C. elegans* with and without hyphae-mediated killing. *Middle:* Survival curve of worms incubated with WT, *hda2 Δ/Δ* and *hda3 Δ/Δ* strains over a 72 hour period. *Right:* The percentage of total worms dead (black bars) and worms killed by hyphae piercing the cuticle (grey bars) after the 72 hour incubation period. Error bars represent the standard deviation of three independent biological replicates.

To test whether the hyphae-defective phenotype observed in *hda2 Δ/Δ* and *hda3 Δ/Δ* isolates is sufficient to impair hyphal growth *in vivo*, we performed killing assays using the nematode *Caenorhabditis elegans* as an infection system (39, 40). These analyses revealed no differences in hyphae formation or percentage of killing between WT, *hda2 Δ/Δ* and *hda3 Δ/Δ* strains (Fig 6D). Therefore, although Hda2 and Hda3 are critical for hyphae-formation in specific media, additional redundant pathways can compensate for the lack of Hda2 and Hda3 in a more complex *in vivo* situation.

### Stability of Hda1 protein is regulated by environmental changes

Comparison of the gene expression profiles in yeast-promoting conditions (YPD 30 °C) and hyphae-promoting conditions (RPMI 37 °C) reveals that deletion of *HDA1* leads to different transcriptional changes in different environments (Fig 7A). In yeast inducing conditions, 2046 genes are significantly differentially expressed upon deletion of Hda1 compared to WT cells, suggesting that Hda1 acts as a global regulator of gene expression (Fig 7A and Dataset S1). In contrast, in hyphae inducing conditions, 507 genes are significantly differentially expressed with hyphal growth being one of the major pathways that is altered (Fig 7B and Dataset S1). These results suggest that Hda1 function is rewired during the yeast to hyphae switch. We hypothesise that this change in function could be due to a differential gene expression profile of *HDA1* in yeast compared to hyphae. While analysis of the RNA-seq dataset reveals that *HDA1, HDA2* and *HDA3* transcription levels are similar across these two conditions, Western analyses clearly demonstrate that levels of Hda1 and Hda3 proteins, but not Hda2, are lower in hyphal cells compared to yeast cells (Fig 7C). Therefore, the functional rewiring of the Hda1 complex upon an environmental change is linked to a reduced expression of this subunit.

**Figure 7.**
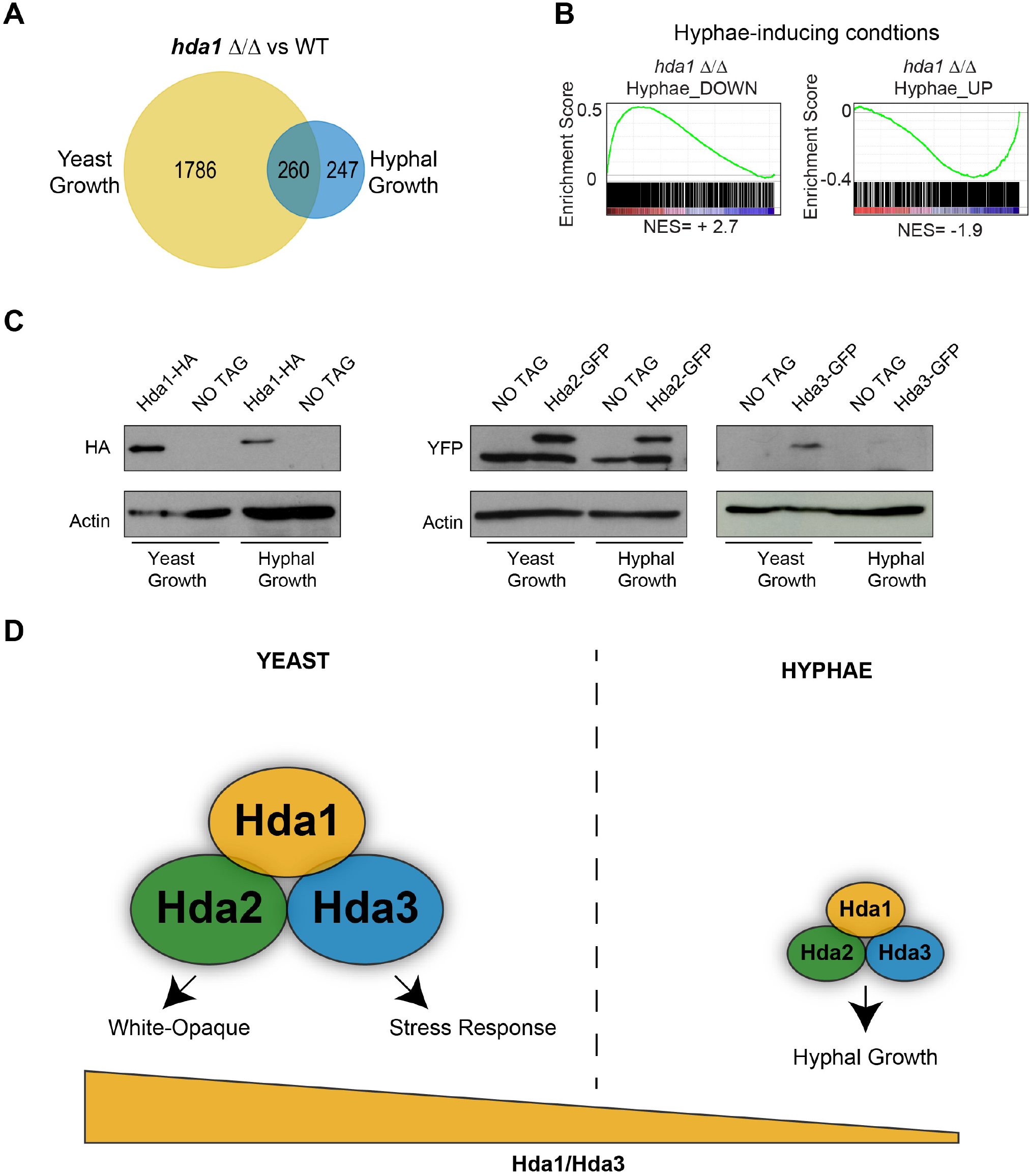
Stability of the Hda1 protein is regulated by environmental changes. **(A)** Venn diagram of genes differentially expressed in *hda1* Δ/Δ relative to WT in yeast and hyphal growth conditions. **(B)** Enrichment plots for genes differentially expressed in *hda1* Δ/Δ relative to genes downregulated (Hyphae_Lee_DN) or upregulated (Hyphae_Lee_up) in hyphae-growth condition in Lee’s media. The x-axis shows genes ranking according to their expression in *hda1A/A* from upregulated (left) to down-regulated (right) genes. Black vertical lines mark individual genes. The cumulative value of the enrichment score (y-axis) is represented by the green line. A positive normalised enrichment score (NES) indicates enrichment in the up-regulated group of genes while a negative NES indicates prevalence of the genes in the down-regulated group. **(C)** HA and GFP Western blots of whole protein extract from strains: Hda1-HA, Hda2-GFP and Hda3-GFP. Actin is shown as a loading control. Cells were grown in yeast (YPD 30 °C) or hyphae-inducing (RPMI 37 °C) conditions. **(D)** Model showing how decreased Hda1 and Hda3 protein levels leading to the functional rewiring of the Hda1 complex in *C. albicans* in different environments. Under yeast growth condition, the Hda1 complex functions as a global regulator of gene expression due to the high level of Hda1 and Hda3. Under hyphae growth conditions, decreased levels of Hda1 and Hda3 leads to specialisation of the Hda1 complex controlling only filamentous growth.

## DISCUSSION

### The role of the Hda1 complex in regulating morphological switches in *C. albicans*

Results presented in this study show that the fungal-specific proteins Hda2 and Hda3 are important regulators of morphological switches in *C. albicans*. We propose that this regulation is mediated via the Hda1 complex. This hypothesis is supported by our observation that Hda1, Hda2 and Hda3 physically interact and by the published results demonstrating that Hda1 controls both the white-opaque and the yeast-hyphae morphological switches (19, 25, 26). Based on the data presented here, it is possible that Hda2 and Hda3 control the activity of the Hda1 complex by different non-mutually exclusive mechanisms. First, Hda2 and Hda3 could mediate Hda1 recruitment to target sites regulating their chromatin acetylation state and their transcriptional activity. Indeed, in *S. cerevisiae*, Hda2 and Hda3 DNA-binding domains are sufficient to bind DNA *in vitro* (32). The structural alignment presented in this study demonstrates that, similar to *S. cerevisiae, C. albicans* Hda2 and Hda3 contain a DNA binding domains with structure resembling the helicase fold found in the SWI2/SNF2-type chromatin-remodeling ATPase (32). We hypothesise that Hda2 and Hda3 could target Hda1 to key genomic locations leading to transcriptional downregulation of associated genes. In support of this hypothesis, we have found that *WOR1*, the master regulator of white-opaque switching, is moderately upregulated in cells deleted for *HDA2* and *HDA3* genes. A similar upregulation is observed in *hda1 Δ/Δ* isolates. We propose that this upregulation is sufficient to activate the Wor1 positive feedback loop promoting white-opaque switching. Indeed, *hda2 Δ/Δ* and hda3 *Δ/Δ* cells undergo white-opaque switching more frequently than WT cells.

Alternatively, interactions between Hda1, Hda2 and Hda3 may induce a conformational change promoting Hda1 deacetylase activity. In support of this hypothesis, it has been established that the *in vitro* catalytic activity of *S. cerevisiae* Hda1 depends on Hda2 and Hda3 (31). This regulation could be critical for controlling deacetylation of non-histone substrates. For example, *C. albicans* Hda1 controls the yeast to hyphae switch by deacetylating a non-histone substrate, the NuA4 component Yng2 (19). We hypothesise that interaction between Hda1, Hda2 and Hda3 causes a conformational change in Hda1 allowing deacetylation of Yng2. Chromatin modifiers are ideal sensors of changing environments as they can respond to external stimuli by rapidly and reversibly changing the transcriptional state of many genes simultaneously. Our results indicate that the activity of the Hda1 complex is rewired in different environmental conditions. While in yeast cells, Hda1 acts as a globular regulator of gene expression, in hyphae-inducing conditions Hda1 function is dedicated to the yeast-hyphae switch. This functional rewiring is accompanied by changes in the protein levels of Hda1 and Hda3. These findings suggest a model through which hyphae-specific function is achieved by diminishing the concentration of key proteins of the Hda1 complex. We did not observe any changes in the RNA levels of *HDA1* and *HDA3* and therefore we hypothesise that translation efficiency and/or stability of these two proteins is differentially regulated in different environments.

Chromatin modifiers are often embedded in multiprotein complexes associated with several non-catalytic subunits that regulate their targeting to substrates or their catalytic activity (24). Our results highlight how Hda2 and Hda3, the non-catalytic subunits of the Hda1 complex, play important roles in regulating morphological switches in response to environmental changes.

### Hda2 and Hda3 as potential targets for anti-fungal therapy

The yeast to hyphae switch is central to *C. albicans* virulence and pathogenesis. Our results signify that while Hda2 and Hda3 promote hyphae formation under specific growth conditions (RPMI and Spider media at 37 °C), they are dispensable for hyphae formation and virulence in the *C. elegans* infection system. We hypothesise that this is due to the presence of redundant pathways that stimulate hyphae formation. Indeed, it has been shown that hyphae induction is a highly orchestrated process integrating several different environmental signals (13). The finding that, *in vivo*, lack of Hda2 and Hda3 does not impair hyphae formation is in agreement with the observation that strains mutant for the *YNG2* gene, the target of Hda1, also do not impair hyphae growth *in vivo* while they are defective for hyphae formation *in vitro* (27).

There is an urgent need to develop new anti-fungal drugs due to emergence of fungal strains resistant to currently available drugs. We propose that Hda2 and Hda3 could be targets for novel antifungal drugs to be used in combination therapy. Combination therapy offers several advantages compared to single-drug therapy. This is because it allows for widening of the spectrum and potency of drug activity and it can also lead to reduction in the dosage of individual agents preventing emergence of antifungal resistance (41). Several features make Hda2 and Hda3 attractive targets for antifungal drug development. First and most importantly, the proteins are present in fungi but absent in humans minimising potential toxicity. Secondly, drugs targeting Hda2 and Hda3 could have a broad spectrum of activity against a variety of human fungal pathogens as Hda2 and Hda3 orthologs are present in other human fungal pathogens, such as *C. glabrata* (CAGL0H01331g and CAGL0G09867g) and *A. fumigatus* (Hda2: Afu5g03390).

KDACs are emerging as promising candidates for drug development and several small molecules inhibiting KDAC activity are currently in clinical trials as potential anti-cancer therapeutics. We propose that inhibition of Hda2 and Hda3 will impair *C. albicans* Hda1 activity but not human KDACs reducing the risk of toxicity.

## MATERIAL AND METHODS

### Growth conditions

Strains are listed in Table 1. Yeast cells were cultured in rich medium (YPD) containing extra adenine (0.1 mg/ml) and extra uridine (0.08 mg/ml), complete SC medium (Formedium) or RPMI medium (Sigma-Aldrich). When indicated, media were supplemented with 30 μg/ml doxycycline. For analysis in stress conditions, YPD agar media was supplemented with 9 nM rapamycin, 2 M sodium chloride, 1 mM SNP with 1 mM SHAM, 7 mM copper sulphate, 300 mM lithium chloride, .01 % sodium dodecyl sulfate (SDS), 15mM caffeine, 1.5 M sorbitol, 18 mM cycloheximide, 1.5 mM cobalt, 25 mM hydroxyurea, 3 % ethanol, 20 μM Calcofluor White, 4.5 mM hydrogen peroxide, 0.75 mM EDTA and 2 μM Cerulenin (42). Cells were grown at 30 °C or 37 °C as indicated.

**Table 1.**
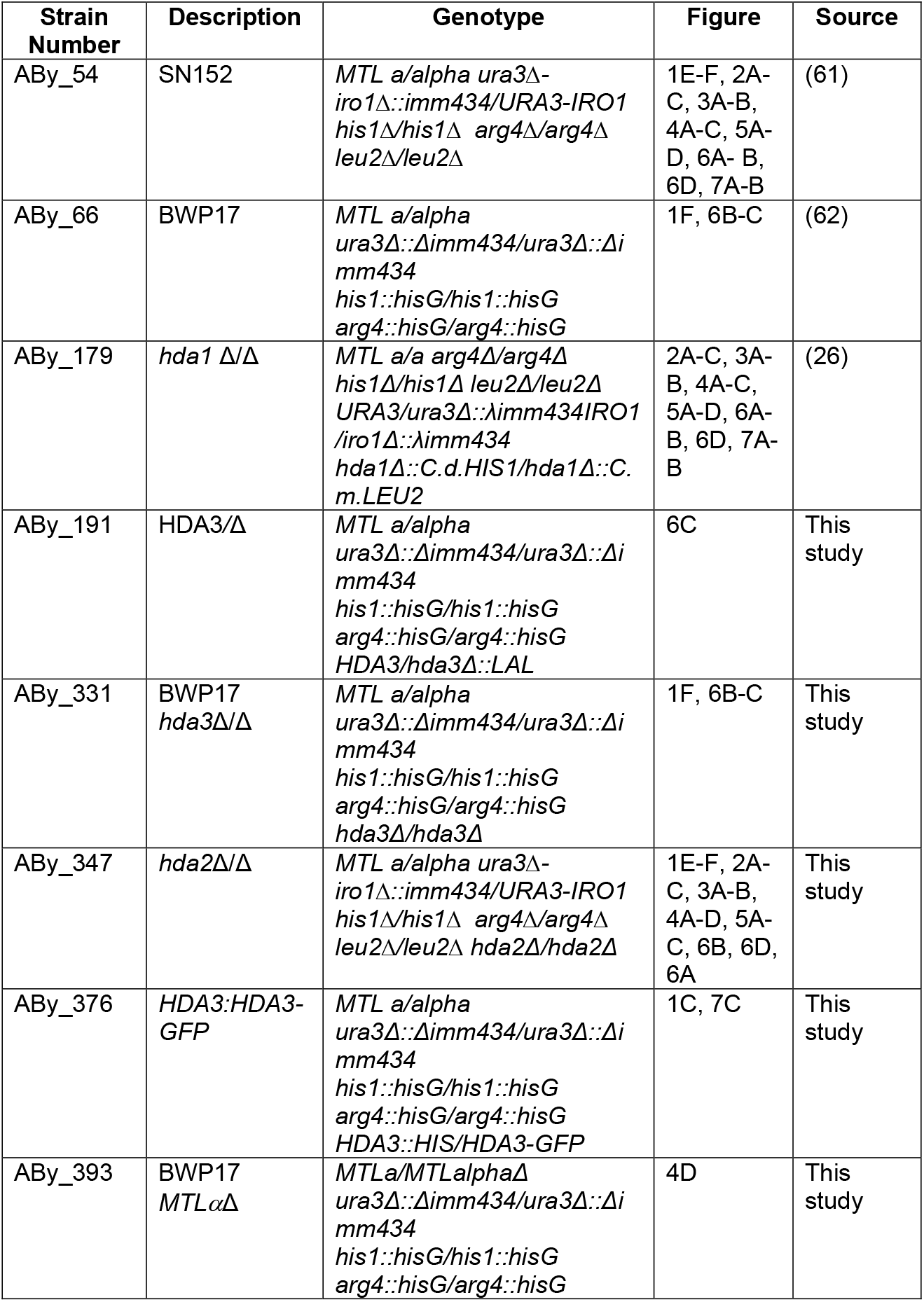
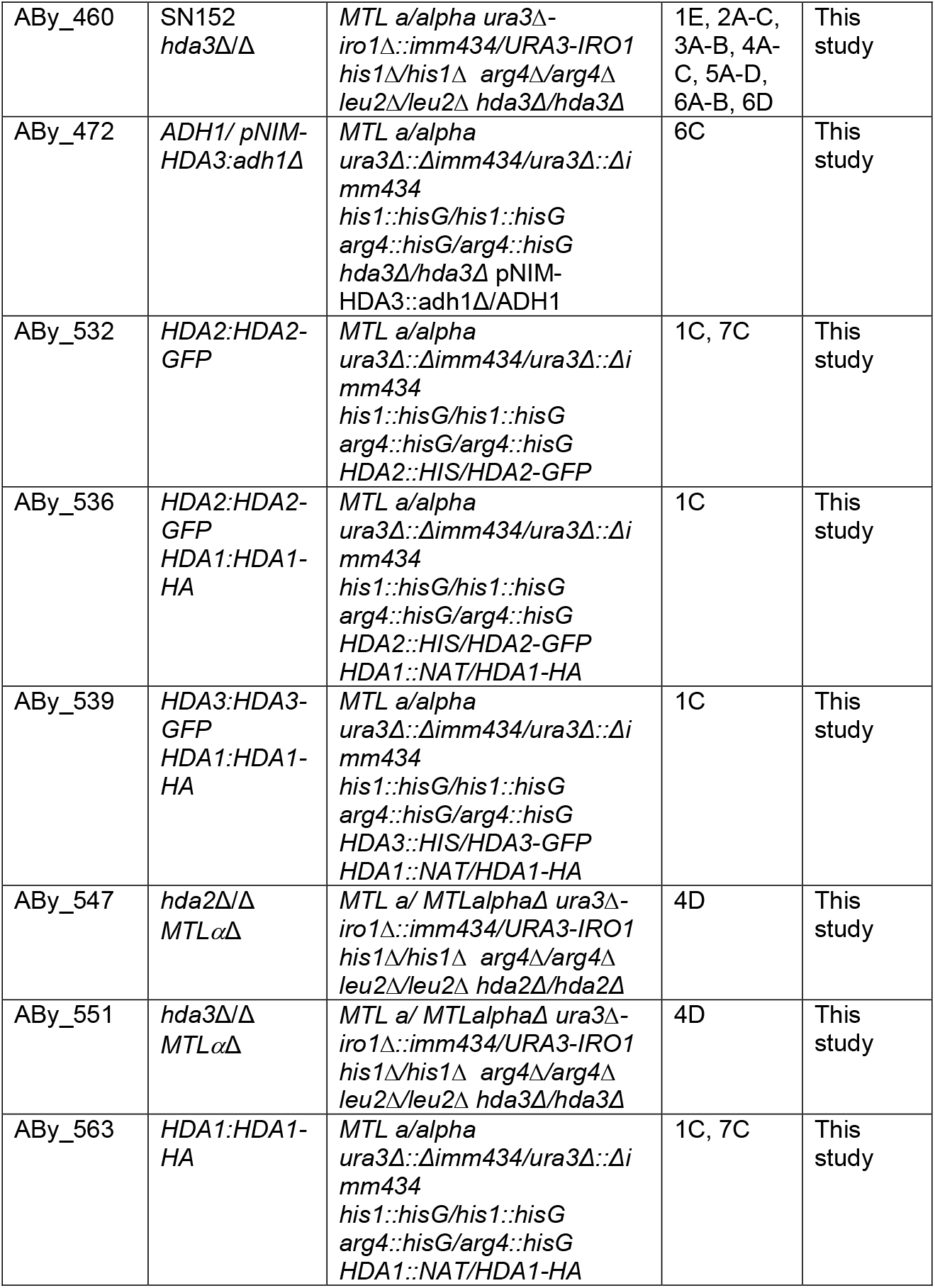
*C.albicans* strains used in this study.

| Strain Number | Description | Genotype | Figure | Source |
| --- | --- | --- | --- | --- |
| ABy_54 | SN152 | *MTL a/alpha ura3Δ-iro1Δ::imm434/URA3-IRO1 his1Δ/his1Δ arg4Δ/arg4Δ leu2Δ/leu2Δ* | 1E-F, 2A-C, 3A-B, 4A-C, 5A-D, 6A- B, 6D, 7A-B | (61) |
| ABy_66 | BWP17 | *MTL a/alpha ura3Δ::Δimm434/ura3Δ::Δimm434 his1::hisG/his1::hisG arg4::hisG/arg4::hisG* | 1F, 6B-C | (62) |
| ABy_179 | *hda1 Δ/Δ* | *MTL a/a arg4Δ/arg4Δ his1Δ/his1Δ leu2Δ/leu2Δ URA3/ura3Δ::λimm434IRO1 /iro1Δ::λimm434 hda1Δ::C.d.HIS1/hda1Δ::C.m.LEU2* | 2A-C, 3A-B, 4A-C, 5A-D, 6A-B, 6D, 7A-B | (26) |
| ABy_191 | HDA3/Δ | *MTL a/alpha ura3Δ::Δimm434/ura3Δ::Δimm434 his1::hisG/his1::hisG arg4::hisG/arg4::hisG HDA3/hda3Δ::LAL* | 6C | This study |
| ABy_331 | BWP17 *hda3Δ/Δ* | *MTL a/alpha ura3Δ::Δimm434/ura3Δ::Δimm434 his1::hisG/his1::hisG arg4::hisG/arg4::hisG hda3Δ/hda3Δ* | 1F, 6B-C | This study |
| ABy_347 | *hda2Δ/Δ* | *MTL a/alpha ura3Δ-iro1Δ::imm434/URA3-IRO1 his1Δ/his1Δ arg4Δ/arg4Δ leu2Δ/leu2Δ hda2Δ/hda2Δ* | 1E-F, 2A-C, 3A-B, 4A-D, 5A-C, 6B, 6D, 6A | This study |
| ABy_376 | *HDA3:HDA3-GFP* | *MTL a/alpha ura3Δ::Δimm434/ura3Δ::Δimm434 his1::hisG/his1::hisG arg4::hisG/arg4::hisG HDA3::HIS/HDA3-GFP* | 1C, 7C | This study |
| ABy_393 | BWP17 *MTLαΔ* | *MTLa/MTLalphaΔ ura3Δ::Δimm434/ura3Δ::Δimm434 his1::hisG/his1::hisG arg4::hisG/arg4::hisG* | 4D | This study |
| ABy_460 | SN152<br>*hda3Δ/Δ* | *MTL a/alpha ura3Δ-iro1Δ::imm434/URA3-IRO1 his1Δ/his1Δ arg4Δ/arg4Δ leu2Δ/leu2Δ hda3Δ/hda3Δ* | 1E, 2A-C, 3A-B, 4A-C, 5A-D, 6A-B, 6D | This study |
| ABy_472 | *ADH1/ pNIM-HDA3:adh1Δ* | *MTL a/alpha ura3Δ::Δimm434/ura3Δ::Δimm434 his1::hisG/his1::hisG arg4::hisG/arg4::hisG hda3Δ/hda3Δ pNIM-HDA3::adh1Δ/ADH1* | 6C | This study |
| ABy_532 | *HDA2:HDA2-GFP* | *MTL a/alpha ura3Δ::Δimm434/ura3Δ::Δimm434 his1::hisG/his1::hisG arg4::hisG/arg4::hisG HDA2::HIS/HDA2-GFP* | 1C, 7C | This study |
| ABy_536 | *HDA2:HDA2-GFP*<br>*HDA1:HDA1-HA* | *MTL a/alpha ura3Δ::Δimm434/ura3Δ::Δimm434 his1::hisG/his1::hisG arg4::hisG/arg4::hisG HDA2::HIS/HDA2-GFP HDA1::NAT/HDA1-HA* | 1C | This study |
| ABy_539 | *HDA3:HDA3-GFP*<br>*HDA1:HDA1-HA* | *MTL a/alpha ura3Δ::Δimm434/ura3Δ::Δimm434 his1::hisG/his1::hisG arg4::hisG/arg4::hisG HDA3::HIS/HDA3-GFP HDA1::NAT/HDA1-HA* | 1C | This study |
| ABy_547 | *hda2Δ/Δ*<br>*MTLαΔ* | *MTL a/ MTLalphaΔ ura3Δ-iro1Δ::imm434/URA3-IRO1 his1Δ/his1Δ arg4Δ/arg4Δ leu2Δ/leu2Δ hda2Δ/hda2Δ* | 4D | This study |
| ABy_551 | *hda3Δ/Δ*<br>*MTLαΔ* | *MTL a/ MTLalphaΔ ura3Δ-iro1Δ::imm434/URA3-IRO1 his1Δ/his1Δ arg4Δ/arg4Δ leu2Δ/leu2Δ hda3Δ/hda3Δ* | 4D | This study |
| ABy_563 | *HDA1:HDA1-HA* | *MTL a/alpha ura3Δ::Δimm434/ura3Δ::Δimm434 his1::hisG/his1::hisG arg4::hisG/arg4::hisG HDA1::NAT/HDA1-HA* | 1C, 7C | This study |

### Plasmid construction

Oligos and plasmids used in this study are listed in Table 2 and 3. Plasmid ABp133 contains the *C. albicans HDA1* gene cloned in frame to a C-terminal HA tag. To generate this plasmid, the full length *HDA1* gene was amplified from plasmid ABp88 (synthesised by GeneArt) with oligos Abo408 and Abo409, containing recognition sites for *XmaI*. The digested PCR product was cloned into plasmid pHA-NAT (ABp17)(43) digested with XmaI. Cloning was confirmed by PCR and Sanger sequencing. *HDA3* was cloned in the pNIM plasmid (ABp111) (44) to generate plasmid ABp177. For this purpose, the full length *HDA3* gene was amplified from plasmid ABp152 (synthesised by GeneArt) with oligos ABo624 and ABo625, containing *XhoI* and BglII restriction sites. ABo625 also supplied a stop codon upstream the restriction site. This PCR fragment was cloned in pNIM digested with *SalI* and *BglII*. Cloning was confirmed by PCR and Sanger sequencing.

**Table 2.**
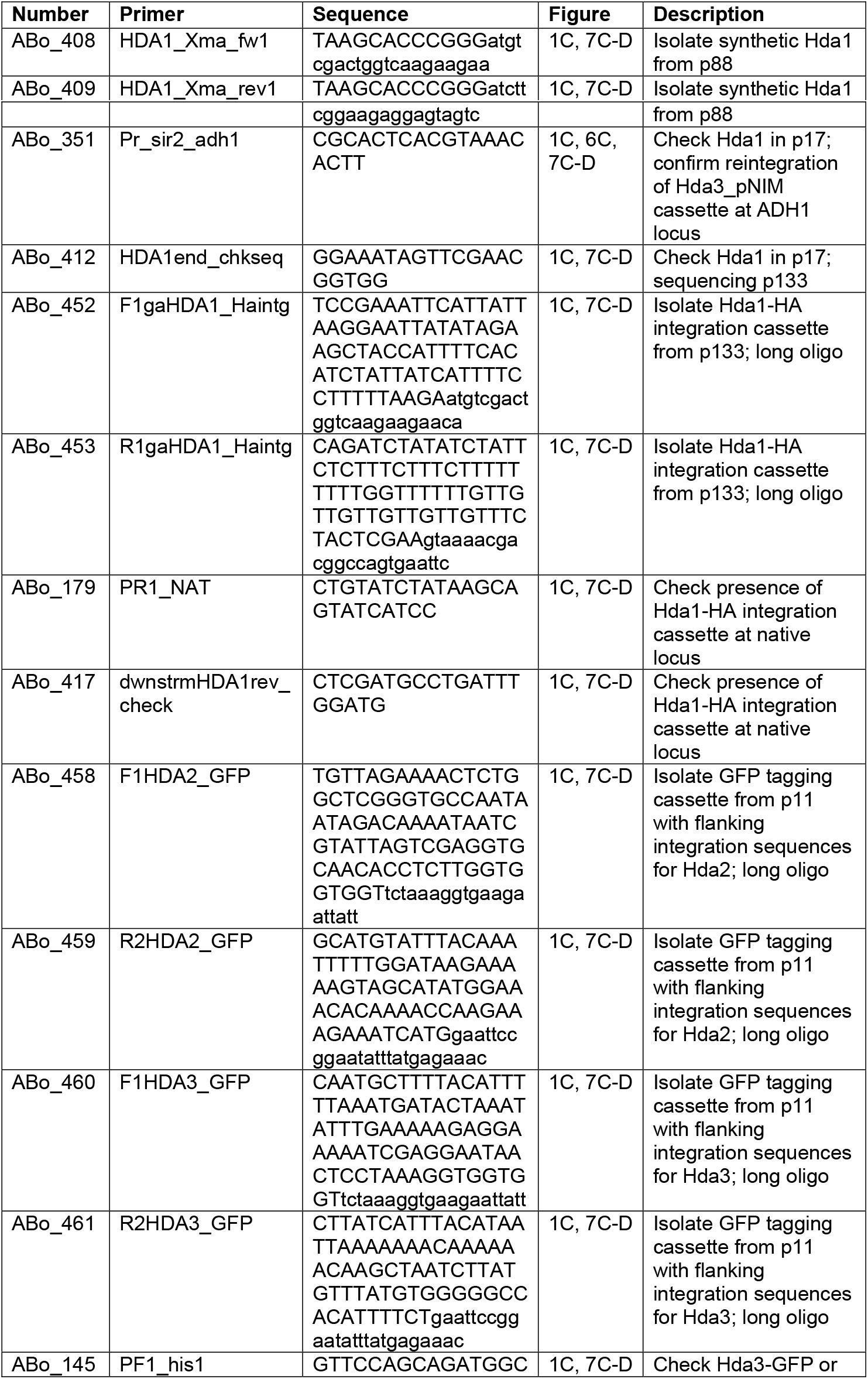
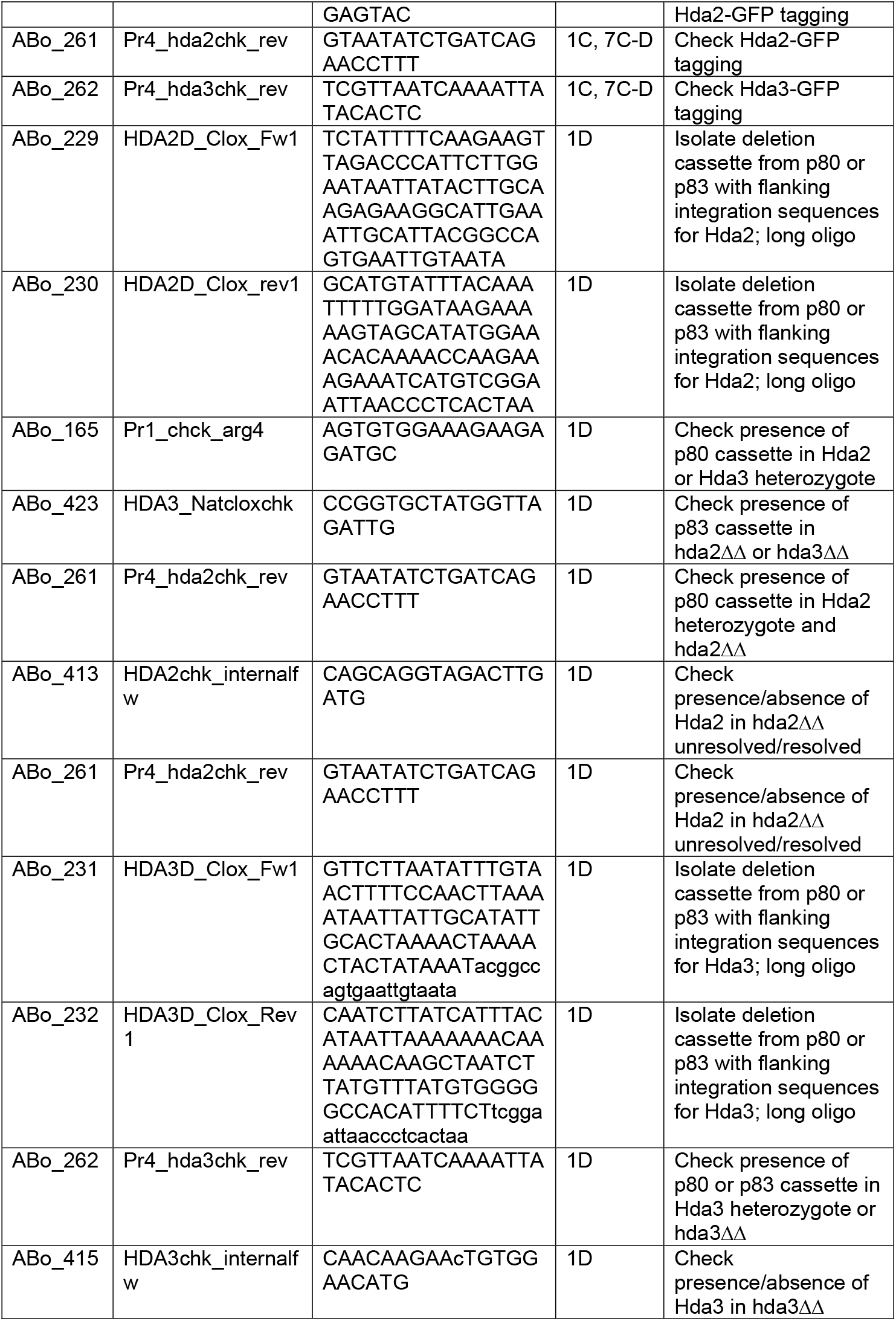
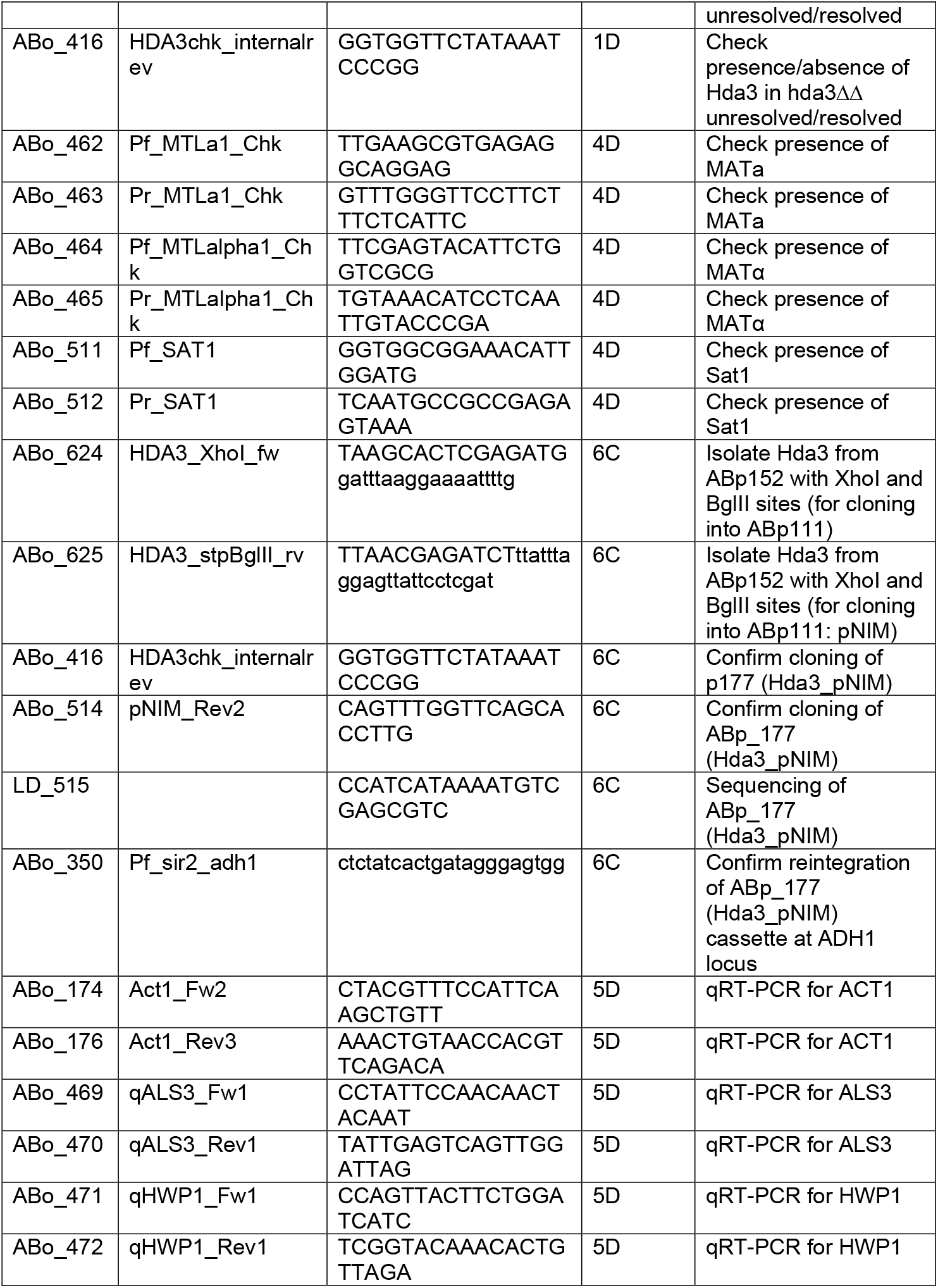
Oligos used in this study.

**Table 3.** Plasmids used in this study.

| ABp_Number | Plasmid | Figure | Description/Source | Source |
| --- | --- | --- | --- | --- |
| ABp_11 | pGFP-His1 | 1C | GFP tagging vector | (45) |
| ABp_17 | pHA_NAT1 | 1C | HA tagging vector | (43) |
| ABp_80 | LAL (loxP-ARG4-loxP) | 1D | *Arg4* substitution products | (33) |
| ABp_83 | NAT1-Clox (loxP-NAT1-MET3p-cre-loxP) | 1D | *Nat* substitution products | (33) |
| ABp_88 | HDA1_synthetic | 1C | Hda1 synthetic | GeneArt |
| ABp_111 | pNIM | 6C | Tetracycline inducible integration to ADH1 locus | (44) |
| ABp_133 | HDA1synthetic_pHA_NAT | 1C | Source of Hda1-HA integration cassette | This study |
| ABp_136 | Sat1 flipper with MTLαKO flanking regions | 4D | Deletion of MATα | Gift from Matthew Anderson Lab, Ohio State |
| ABp_152 | HDA3 synthetic | 6C | HA-HDA3 synthetic: 6x CUG->TCA | GeneArt |
| ABp_177 | HDA3-pNIM | 6C | Tetracycline inducible integration of Hda3 to ADH1 locus | This study |

### Construction of *C. albicans* mutants

Deletions of *HDA2* and *HDA3* were generated in the SN152 or BWP17 background using the *Clox* system for gene disruption (33) using long-oligos PCR, the LAL (loxP-ARG4-loxP) and NAT1-Clox (loxP-NAT1-MET3p-cre-loxP) plasmids as templates. During all selections for *Clox* transformants media was supplemented with 2.5 mM methionine and 2.5 mM cysteine to repress the *MET3* promoter and minimize Cre-loxP mediated recombination. Nourseothricin resistant (Nou^R^) transformants were selected using 200 μg/ml nourseothricin (Melford). *HDA2* and *HDA3* gene deletions were confirmed by PCR and markers were resolved by allowing Cre expression in medium lacking methionine and cysteine as previously described (33).

C-terminal GFP tagging of *HDA2* and *HDA3* genes at the endogenous loci was performed by long oligos PCR using plasmid pGFP-His1 (ABp11) as a template (45). C-terminal HA tagging of Hda1 was performed by long oligos PCR using plasmid ABp133 as a template and transformation into BWP17 (ABy215), HDA2-GFP (ABy532) and HDA3-GFP (ABy376).

For rescue experiments, *HDA3* re-integrated at the *ADH* locus by digesting ABp177 with *KpnI* and SacII and transforming the product into the *hda3 Δ/Δ* (Aby33) deletion strain. Correct integration was confirmed by colony PCR. Transformations of *C. albicans* strains were performed by electroporation using the protocol described in (46) or by lithium acetate transformation (47) of competent cells (48) with a *S. cerevisiae* method adapted to *C. albicans* (49).

### Hyphal growth induction and quantification

Cultures were grown overnight (~16 hours) in 5 ml liquid YPD at 30 °C. Overnight cultures were diluted 1:100 in YPD media and grown ~5 hours at 30 °C. 50 – 100 cells were plated to RPMI or Spider agar and incubated for ~6 days at 37 °C. For doxycycline induction, the drug was added to the media at 30 μg/ml in 20 ml plates.

Phenotypes were documented by imaging with a Leica MZFLIII microscope at 10x to 16x magnification A minimum of 150 colonies were counted for each strain. For liquid filamentation assays, overnight cultures were grown throughout the day and from this stationary culture, 1 mL was harvested and washed with distilled water before resuspending in 10 mL pre-warmed final media (YPD (control) or RPMI). These cultures were grown at 37 °C (control at 30 °C) for 15-17 hours before imaging on an Olympus IX81 inverted microscope at 60x magnification. Samples were evaluated on four separate days and a minimum of 400 cells were counted per sample.

### White-opaque switching essay

Quantitative switching assays were performed as previously described (12) with modifications. Briefly, strains were streaked from frozen stocks on YPD plates and grown at 30 °C for 2 days. Single colonies were picked, resuspended in sterile water and spread onto synthetic complete agar containing 5 μg/ml Phloxine B (Sigma-Aldrich). Formation of opaque colonies or sectors was scored after 9 days. Experiments were done in biological duplicate or triplicate on at least three separate days. A minimum of 400 total colonies were counted for each strain.

The ggplot package in R studio was used to construct violin plots. Unpaired t-tests were performed to test for significant differences between wild-type and mutant strains.

### Structural modelling

The model of Hda3 DBD and CCS domain was produced using Phyre2 in intensive mode (50) and visualised using PyMol (The PyMOL Molecular Graphics System, Version 1.8 Schrödinger, LLC.)

### Whole cell extracts

Preparation of whole cell extracts was performed as described (51). Briefly, overnight YPD cultures were diluted in YPD or RPMI and grown to OD_600_ = ~0.8 at 30 °C. Cells were harvested, resuspended in 200 μl lysis buffer (0.1 M NaOH, 0.05 M EDTA, 2% SDS, 2% β-mercaptoethanol) and heated for 10 minutes at 95 °C before adding 5 μl of 4 M Acetic acid and incubating for further 10 minutes at 90 °C. Extracts were mixed with 50 μl loading buffer (0.25 M Tris-HCl pH 6.8, 50 % Glycerol, 0.05 % Bromophenol Blue), incubated at 96 °C for 5 minutes and centrifuged at 13000 rpm for 5 minutes. Supernatants were collected and analysed by SDS-PAGE and Western blot analyses.

### Immunoprecipitation

Immunoprecipitation was performed as described (52) with modifications. 1 L YPD cultures (OD_600_ = 1) were harvested at 4000 rpm. Cell were washed 3 times in cold water and resuspended in 1/5th volume (water/cells). Cell pellets were ground in liquid nitrogen using a mortar and pestle for 30 minutes and resuspended in 10 ml of cold lysis buffer (50 mM HEPES-NaOH pH 7.5, 150 mM NaCl, 5 mM EDTA, 0.1 % NP-40, 5 mM DTT, 1x Roche EDTA-free protease inhibitor cocktail and 0.2 mM PMSF added fresh before use). The cells were solubilized for 30 minutes with rotation at 4 °C. Following centrifugation, supernatant samples were incubated with 4 μl of magnetic beads pre-coupled with anti-HA antibody (Sigma-Aldrich) for 2 hours with rotation at 4 °C. Beads were washed four times in lysis buffer and analysed by SDS-PAGE and Western blot.

### Antibodies information

The following antibodies were used for Western analyses. Anti-HA antibody (#11666606001, Sigma-Aldrich) diluted at 1:1000, anti-GFP antibody (Roche #1184460001) diluted at 1:5000, anti-actin (Cooper Lab, Washington University, St. Louis, Mo., USA) diluted at 1:5000.

### RNA extraction

Overnight YPD cultures were diluted in YPD and grown to OD_600_ = ~0.8. Cells were pelleted, washed once with sterile water and resuspended in pre-warmed YPD (30 °C) or RPMI (37 °C) for 90 minutes. RNA extraction was performed using a yeast RNA extraction kit (E.Z.N.A. Isolation Kit RNA Yeast, Omega Bio-Tek) following the manufacturer’s instructions with the following modifications: (1) 30 °C incubation in SE Buffer/2-mercaptoethanol/lyticase solution time was increased to 90 minutes; (2) lysis was performed with bead mill at top speed for 30 minutes at 4°C. RNA was treated with DNAse I and RNA quality was checked by electrophoresis under denaturing conditions in 1% agarose, 1× HEPES, 6% Formaldehyde (Sigma-Aldrich). RNA concentration was measured using a NanoDrop ND-1000 Spectrophotometer. cDNA synthesis was performed using qPCRBIO cDNA Synthesis Kit (PCR Biosystems) following manufacturer’s instructions.

### High-throughput RNA sequencing

Strand-specific cDNA Illumina Barcoded Libraries were generated from 1 μg of total RNA extracted from WT, *hda1 Δ/Δ*, *hda2 Δ/Δ* and *hda3 Δ/Δ* and sequenced with an Illumina iSeq2000 platform. Illumina library preparation and deep-sequencing was performed by the Genomics Core Facility at EMBL (Heidelberg, Germany). RNA sequencing was performed in duplicates. Raw reads were analysed following the RNA deep sequencing analysis pipeline using the Galaxy platform (https://usegalaxy.org/). Downstream analysis of differential expressed genes was performed with R Studio (https://www.rstudio.com/). Scatter Plot matrices, using Pearson correlation coefficients, were generated with the ggplot package. Heatmaps were generated with the pheatmap package and Pearson correlation for clustering. Heatmaps show the log2 fold changes of differentially expressed genes in *hda1 Δ/Δ, hda2 Δ/Δ* or *hda3 Δ/Δ* compared to wild-type expression. Venn diagrams were generated using the FunRich programme(53). RNA sequencing data are deposited into ArrayExpress (accession number: E-MTAB-6920).

### qRT-PCR

qRT-PCR was performed in the presence of SYBR Green (Bio-Rad) on a Bio-Rad CFXConnect Real-Time System. Data was analysed with Bio-Rad CFX Manager 3.1 software and Microsoft Excel. Enrichment was calculated over actin. Histograms represent data from three biological replicates. Error bars: standard deviation of three biological replicates generated from 3 independent cultures of the same strain.

### Functional analysis and modelling of transcriptional profiles

Gene set enrichment analysis (GSEA) (34, 54) was performed using the GSEA PreRanked tool to determine whether a ranked gene list exhibited statistically significant bias in their distribution within defined gene sets (55) and (*Sellam et al*, personal communication). The weighted enrichment statistics were calculated on 10512 gene sets each containing 5-1000 genes, and the false discovery rate (FDR) was calculated from 1000 permutations. Selected results graphs are shown. Since enrichment profiles can exhibit correlations with hundreds of overlapping gene sets, Cytoscape 3.6 (http://www.cytoscape.org) (56) and the Enrichment Map Pipeline Collection plug-ins (http://apps.cytoscape.org/apps/enrichmentmappipelinecollection) were used to further organise and visualise the GSEA. Enrichment maps were calculated using default parameters.

### *C. albicans-C. elegans* pathogenesis assay

The *C. elegans glp-4/sek-1* strain was used as described previously (39, 57). Briefly, *C. elegans* were propagated on nematode growth medium (NGM) on lawns of *E. coli* OP50. The *C. albicans-C. elegans* pathogenesis assay was performed as previously described (40). Briefly, 100 μl of *C. albicans* cells from an overnight culture were spread into a square lawn on a 10 cm plate containing brain-heart infusion (BHI) agar and kanamycin (45 μg/ml). These were incubated for approximately 20 hours at 30 °C. Synchronized adult *C. elegans glp-4/sek-1* nematodes grown at 25°C were carefully washed from NGM plates using sterile M9 buffer. Approximately 100 to 200 washed animals were then added to the centre of the *C. albicans* lawns and the plates were incubated at 25°C for 4 hours. Worms were then carefully washed into a 15 ml tube using 10 ml of sterile M9, taking care to minimize the transfer of yeast. Worms were washed four or five times with sterile M9. 30-40 worms were then transferred into three wells of a six-well tissue culture plate (Corning, Inc.) containing 2 ml of liquid medium (80% M9, 20% BHI) and kanamycin (45 μg/ml). Worms were scored daily into one of three categories: alive, dead with hyphae piercing the cuticle, and dead without hyphae piercing the cuticle. Worms were considered to be dead if they did not move in response to mechanical stimulation with a pick. Dead worms were removed from the assay. *C. elegans* survival was examined by using the Kaplan-Meier method and differences were determined by using the log-rank test using OASIS 2 tool (58). Differences in the number of worms with *C. albicans* hyphal formation were determined by using a one-way ANOVA with Dunnett’s test for multiple comparisons. The *C. elegans* pathogenesis assay presented here is an average of three independent biologic replicates. A p-value of <0.05 was considered statistically significant.

## ACKNOWLEDGMENTS

We thank GeneCore at EMBL for performing Illumina library preparation and sequencing. We thank Dr Sellam (Biotechnology Research Institute, National Research Council of Canada) and Dr Traven (Monash University) for help with the GSEA analysis. We are grateful to Dr Anderson (The Ohio State University) for the plasmid allowing deletion of mating type cassette and Prod Alistair Brown (University of Aberdeen) for the Clox system. We thank members of the Kent Fungal Group for useful discussions and critical reading of the manuscript.

